# The Effects of DMSO Cryopreservation on the Biomechanics and Histology of Human Cerebrovascular Tissue

**DOI:** 10.1101/2025.09.29.679363

**Authors:** Maximos P McCune, Mark A Davison, Nishanth Thiyagarajah, Daniel E Thomeer, William Albabish, Melissa Owusu-Ansah, Nina Z Moore

## Abstract

**Introduction:** Tissue preservation techniques, chiefly cryopreservation, have been demonstrated to alter vascular histology and tissue biomechanics via rapid osmotic change—resulting in collagen fiber rearrangement and internal elastic laminae (IEL) microfractures; however, this has not yet been evaluated in cerebrovascular tissue. As such, we sought to measure the effectiveness of a canonical cryopreservation strategy on the thickness and continuity of human cerebrovascular tissue. With the recent rise in biomechanical analyses of cerebrovascular tissue for the design of novel treatments and optimization of surgical strategies, the importance of designing models with accurate tissue proxies is paramount.

**Methods:** Fresh, human cerebrovascular tissue was obtained through the Cleveland Clinic institutional cadaver donation program. Donors with prior craniotomy, intracranial malignancy, or history of cerebrovascular disease were excluded. Cadaveric tissue dissections were completed within fourteen days of patient expiration and sectioned into four specimens. The 164 tissue samples obtained from three donors were then randomized into one of the following experimental conditions: 10% formalin (control), distilled water (dH2O), dimethyl sulfoxide (DMSO), or −80 °C DMSO cryopreservation. Specimens were then processed into paraffin-embedded sections and treated with Movat pentachrome staining. Vessel layers were measured by two blinded evaluators and discontinuities in internal elastin lamina were tallied.

**Results:** We found that DMSO cryopreservation failed to consistently provide a protective effect to cerebrovascular specimens. Tissue stored via this method was reported to occasionally swell in specific vessel tunics of select vessel territories compared to formalin controls. We also observed an increase in the number of transverse elastin breaks with DMSO cryopreservation.

**Conclusions:** This data demonstrates that conventional tissue preservation methods may fail to preserve layer thicknesses between some vessels and alter biomechanical properties for future testing. Further, with more frequent elastin fractures in the cryopreservation group, recoilability of preserved vessels may vary from *in vivo* counterparts.

**GRAPHICAL ABSTRACT:** 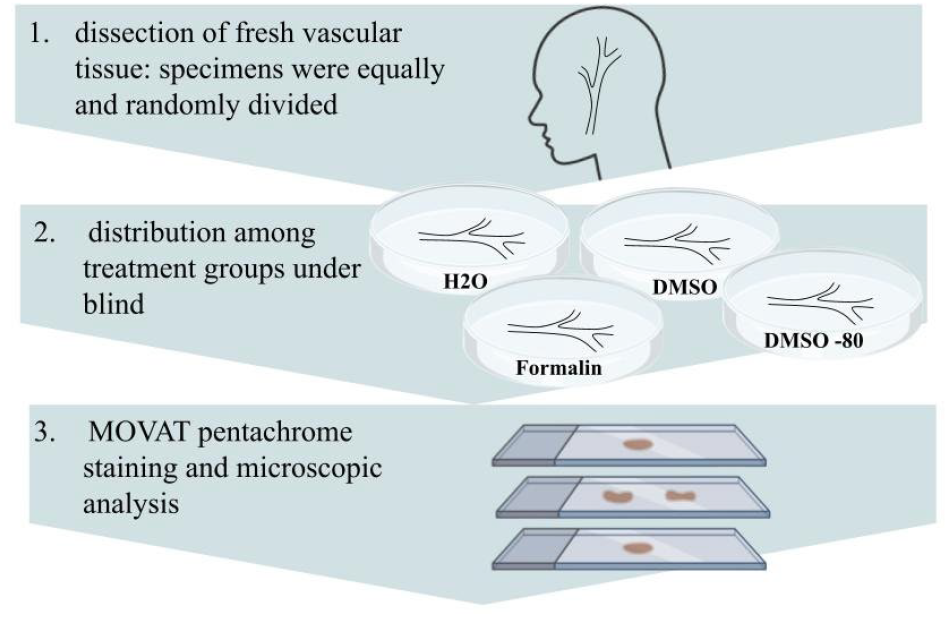

## INTRODUCTION

Cerebrovascular disease is the leading contributor to mortality worldwide, with stroke as the second-most prominent cause of death globally and a greatest source of disability [1,2]. Over 6.5 million deaths and more than 100 million prevalent cases worldwide were attributed to stroke in 2019—rising to 7.3 million deaths in 2021 [1,3]. This steady increase in stroke incidence called for a compensatory surge in pharmacological and neurosurgical interventions.

In recent years, the number of patients receiving both intravenous thrombolysis and mechanical thrombectomy has increased substantially [4,5]. Thrombectomy specifically has seen a drastic rise in popularity, with the window for implementation increasing to 24 hours after last known well [6]. Likewise, potentially life-threatening cerebrovascular anomalies, such as arteriovenous malformation (AVM), are undergoing a change in treatment thresholds. AVMs, a vascular pathology with an incidence rate of 10-15 per 100,000 persons [7–9], are characterized as an abnormal convolution of arteries and veins that contribute substantially to the etiology of intracranial hemorrhage in young adults. AVMs present an annual rupture risk of approximately 2-4% [10]; however, the characteristics used to tailor preclusive rupture interventions are based largely on retrospective analyses [10–12]. Surgical resection of AVMs has declined since the 2014 “A Randomized Trial of Unruptured Brain Arteriovenous Malformations” (ARUBA) found that medical interventions lowered mortality rates more than surgery for those harboring unruptured AVMs, specifically [13]. Between 2014 and 2017, the likelihood of intervention decreased from 28.1% to 22.3%, while admissions due to AVM rupture increased by 3.9% over the same time period [14]. It is possible that the inverse relationship between surgical intervention rates and AVM bleeds is related to the inherent risk of resecting an unruptured AVM.

As such, more detailed studies of vessel biomechanics are needed to optimize neurovascular surgery—to both increase the efficiency of routine thrombectomy for large vessel occlusions and allow for the safe resection of complex vascular anomalies. Such investigation of cerebrovascular biomechanics requires accurate proxies of tissue structure. The unique biomechanical properties of the various intracranial vessel layers determine compliance, elasticity, and failure modes, which in turn influence the growth and tolerance of vascular pathology [15]. Accurate characterization of these profiles is therefore essential for tailoring interventions and optimizing surgical outcomes.

Intracranial arteries are composed of three principal tunics: the intima (consisting of the endothelium and subendothelial matrix), the media (which contains smooth muscle cells and concentric elastic lamellae), and the adventitia (comprised of collagen-rich connective tissue and vasa vasorum). The internal elastic lamina (IEL) lies within the innermost tunic and provides vascular elasticity, enabling vessel recoil following diastole and providing structural reinforcement [16]. Thiyagarajah *et al*. demonstrated that the IEL is significantly thicker in the anterior circulation vessels than in the posterior (6.01 µm vs 4.4 µm, p < 0.05) [17], suggesting that certain regional disparities exist that may constitute differences in vascular pathology. Detecting such differences in vessel morphology requires well-preserved specimens and careful consideration of dissection technique. Accurate characterization of vessel layer properties necessitates preservation methods that maintain tissue architecture.

Vascular tissue has traditionally been preserved via chemical fixation or cryopreservation. Cryopreservation strategies, including slow freezing and vitrification, use agents such as dimethyl sulfoxide (DMSO) to suppress ice crystal formation. As a means of practical convenience, vascular tissue is typically cryopreserved to halt tissue degradation that may occur during the period between harvesting and testing of biomechanical properties [18]—with DMSO being among the more common storage mediums. DMSO functions to preserve cellular membrane flexibility by preventing intracellular ice nucleation [19]. Spectrographic analysis found that DMSO disrupts hydrogen bond networks and halts the propagation of ice crystal formation within tissue [20]. Additionally, cryoprotectants function to decrease the temperature at which tissue dehydration occurs by increasing subzero membrane permeability [21].

Despite this apparent protective effect, DMSO can also impose hyperosmotic stress during the freezing and thawing phases of tissue preservation, damaging cellular matrix components. Prior works have noted post-thaw changes in vascular tissue after DMSO cryopreservation; this was postulated to be the result of thermal stresses during the thawing/reheating processes [22]. Prior investigations of larger caliber vessels underscore these risks. A study of porcine iliac arteries found denudation and significant fracture in light of DMSO cryopreservation following rapid thawing [23]. Further, the biomechanical properties of vascular tissue was shown to change in the setting of cryopreservation [24] [25]. To evaluate whether canonical DMSO cryopreservation preserves cerebrovascular microarchitecture, cadaveric intracranial vessels were subjected to standard DMSO protocols and observed for anatomical change.

## METHODS

### Tissue Procurement & Preservation

Cerebrovascular tissue was collected from three human cephalids provided by the Cleveland Clinic cadaver donation program. Donors with prior craniotomy, intracranial malignancy, or history of cerebrovascular disease were excluded. Vessels were extracted within 14 days of donor death and sectioned into 4 specimens. All 3 donors were male, with an average age at the time of death of 72.67 years (Table 1). Prior analysis suggests that elastin area remains consistent with age [26], which remedies some concern for the elevated age of donors in this study. A total 41 vessels of interest were harvested across the three cadaveric dissections, which included the following: communicating segment of the Internal Carotid Artery (ICA), Anterior Cerebral Artery (ACA. A1, A2), Middle Cerebral Artery (MCA. M1, M2, M3, M4), Posterior Cerebral Artery (PCA. P1, P2), Superior Cerebellar Artery (SCA), Pericallosal Artery, and Basilar Artery (Table 2).

**Table 1.**
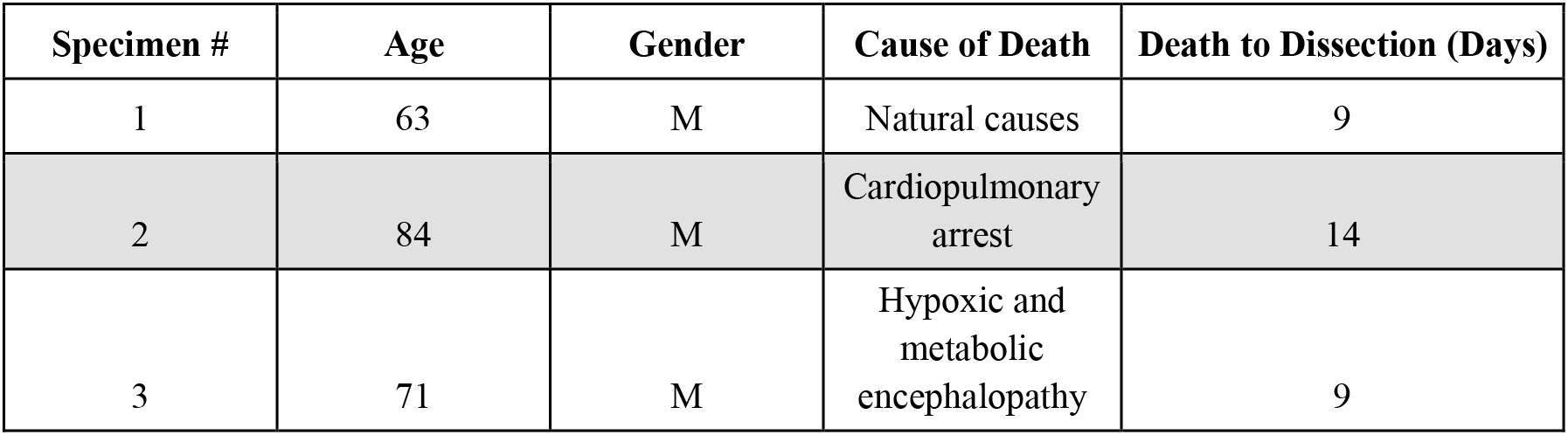
Cadaveric specimen demographics.

| Specimen # | Age | Gender | Cause of Death | Death to Dissection (Days) |
| --- | --- | --- | --- | --- |
| 1 | 63 | M | Natural causes | 9 |
| 2 | 84 | M | Cardiopulmonary arrest | 14 |
| 3 | 71 | M | Hypoxic and metabolic encephalopathy | 9 |

**Table 2.**

| Territory Groups | Vessel | Count |
| --- | --- | --- |
| Anterior Circulation | Internal Carotid Artery | 2 |
|  | Anterior Cerebral Artery | 9 |
|  | Middle Cerebral Artery | 20 |
|  | Anterior Temporal Artery | 1 |
|  | Pericallosal Artery | 2 |
| Posterior Circulation | Superior Cerebellar Artery | 2 |
|  | Posterior Cerebral Artery | 4 |
|  | Basilar Artery | 1 |

Vessel lumens were rinsed post-extraction with distilled water (dH_2_O) to clear the tissue of macroscopic calcifications and loose atherosclerotic buildup. Vessels for which the gross morphology was damaged by plaque or macerated during tissue procurement were excluded. Each section was randomized into one of four experimental groups: 10% formalin (control), distilled water, 10% DMSO, or 10% DMSO with −80 °C freeze to infer within-group comparisons. dH_2_O was included as an additional negative control, as it was used during both the vessel collection process and to dilute the formalin used for room temperature (RT) storage. Specimens treated with dH_2_O and DMSO were placed into 10% formalin after twenty four-hour refrigeration, while −80 °C DMSO samples were frozen then meticulously thawed after fourteen days. Following experimental exposure to one of the four treatment groups, vessel specimens were placed into 10% formalin and stored at room temperature until tissue processing and staining.

### Histological Preparations

5-µm sections were then taken from each specimen and fixed in paraffin. The sections were then mounted and stained with Movat pentachrome for visualization of elastin, adventitia, and smooth muscle vessel layers. Each section was examined for gross malformation or staining error before detailed analysis. Slides were imaged via Akoya Vectra Polaris high-resolution slide scanner at 20x magnification and assigned a unique specimen ID.

### Tissue Scoring

Two trained scorers—both blinded to the vessel territory and experimental condition— independently evaluated each specimen, taking measurements of individual layer thicknesses and elastin quality. The diameter of each layer and the number of elastic breaks due to experimental manipulation were tallied. Large transverse fractures of elastin resulting from tissue manipulation during the sectioning or staining processes were not included. Each reviewer measured elastin, smooth muscle, adventitia, and total wall thickness at four distinct locations about the perimeter of each vessel. Measurements were averaged to provide a layer thickness value for each specimen. All tissue scoring was accomplished in Fiji, version 2.17.0 [27].

### Mechanical Inflation Testing

To measure the effect of cryopreservation on cerebrovascular biomechanics, we biaxially strained a series of right M3 vessels via inflation testing. Each vessel was pressurized to 1.5 and 2.5 PSI. Laser micrometers were used to measure vessel distension once pressurized at a sampling rate of 10 Hz. Vessels underwent near-instantaneous pressurization and were allowed to reach steady state.

### Statistical Analysis

To ensure inter-rater reliability, Bland-Altman analyses were performed for each set of layer measurements between the two independent reviewers. Limits of agreement were set based on a 95% confidence interval. We report mean differences of 0.34 ± 2.82 (elastin), 9.98 ± 27.66 (smooth muscle), 5.6 ± 27.98 (adventitia), and 9.56 ± 36.81 (total wall) resulting in an average absolute bias across all layers of 6.37. Average differences between matched comparisons of vessel measurements fell within 4.62-10.2% of the average tissue thickness, which we deemed acceptable for the sake of comparisons put forth in this study (Table 3). Data from a single reviewer was compiled for descriptive statistics; despite showing favorable inter-rater reliability, measurements from both reviewers were not pooled to limit discrepancies between vessel measurements and avoid near-duplications of data. Relegating statistical calculations to a single reviewer removes the need to consider the few overtly disparate matched comparisons beyond our limits of agreement (Figure 1).

**Table 3.**

| Layer | Mean Layer Thickness ± SD ($\mu\text{m}$) | \|Bias\| ± SD ($\mu\text{m}$) | LLoA | ULoA | Standard Error of LoAs | Bias % of Mean Layer Thickness (%) |
| --- | --- | --- | --- | --- | --- | --- |
| Elastin | 7.41 ± 2.58 | 0.34 ± 2.82 | -5.87 | 5.19 | 0.2 | 4.59 |
| Smooth Muscle | 97.92 ± 36.95 | 9.98 ± 27.66 | -64.21 | 44.24 | 1.94 | 10.19 |
| Adventita | 75.52 ± 28.38 | 5.6 ± 27.98 | -49.25 | 60.44 | 1.96 | 7.42 |
| Total Vessel Thickness | 189.86 ± 50.05 | 9.56 ± 36.81 | -81.71 | 62.59 | 2.58 | 5.04 |

**Figure 1.**
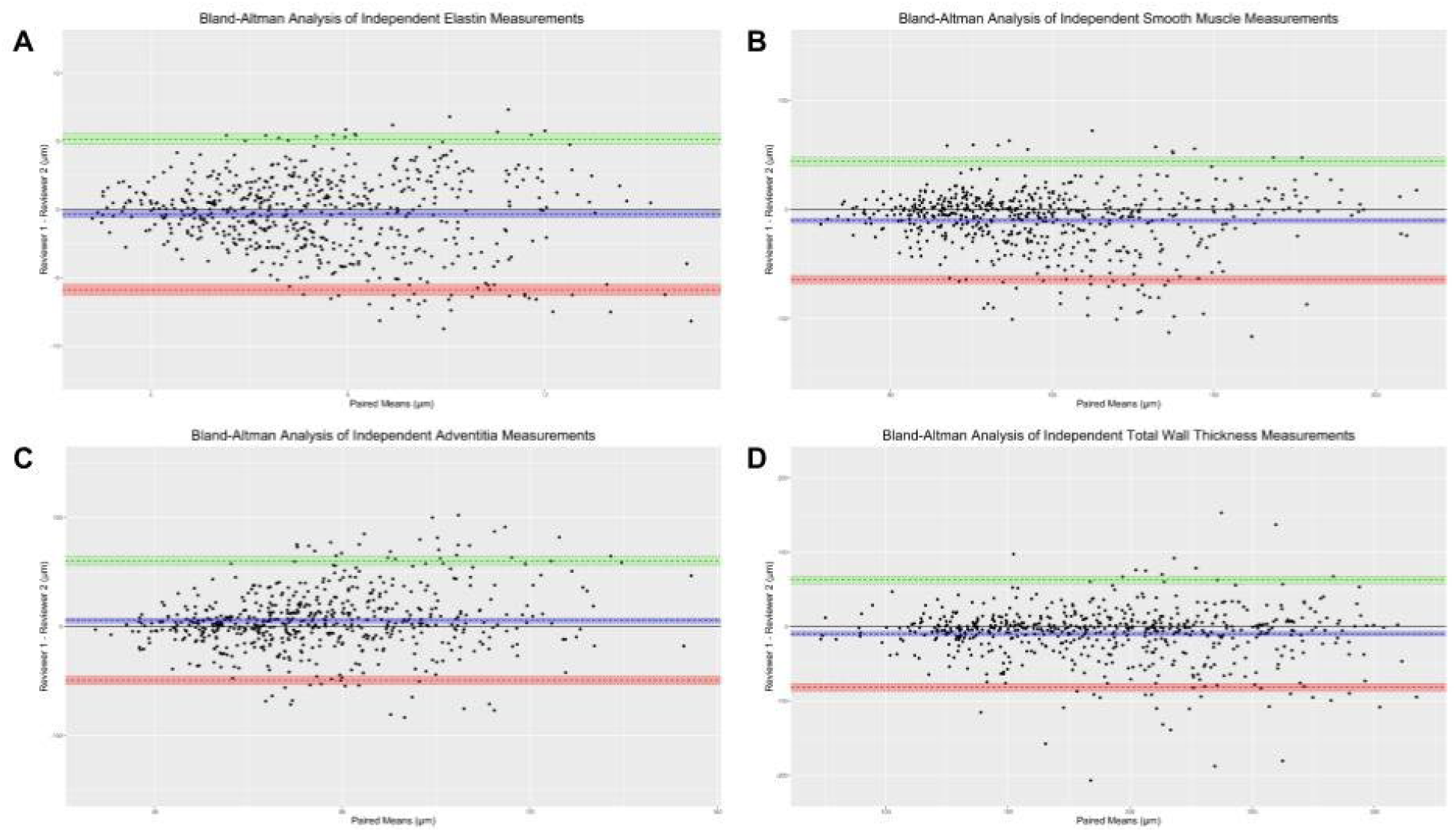
Bland-Altman analyses of elastin (A), smooth muscle (B), Adventitia (C), and total vessel wall (D) thicknesses are represented herein. The solid line denotes the zero bias axis, with dashed lines representing bias (blue), upper limit of agreement (green), or lower limit of agreement (red). Shaded regions mark the range of upper and lower 95% confidence intervals.

Specimens with measurements > 1.5 times the interquartile range of their respective layer and vessel territory groups were considered outliers and removed, leaving a total of 41 specimens for analysis (Table 2). Vessel measurements reported herein are continuous and presented as means and standard deviations. Statistical analyses were two-sided and p-values < 0.05 were considered significant. All statistical calculations were performed using R version 4.4.0 (R Foundation, Vienna, Austria).

## RESULTS

### Effects of DMSO on Gross Vessel Morphology

Upon microscopic examination, the gross vessel morphology of quantified vessel layers was intact. Figure 2 depicts a model vessel scored in this study, with contiguous layers and an intact IEL.

**Figure 2.**
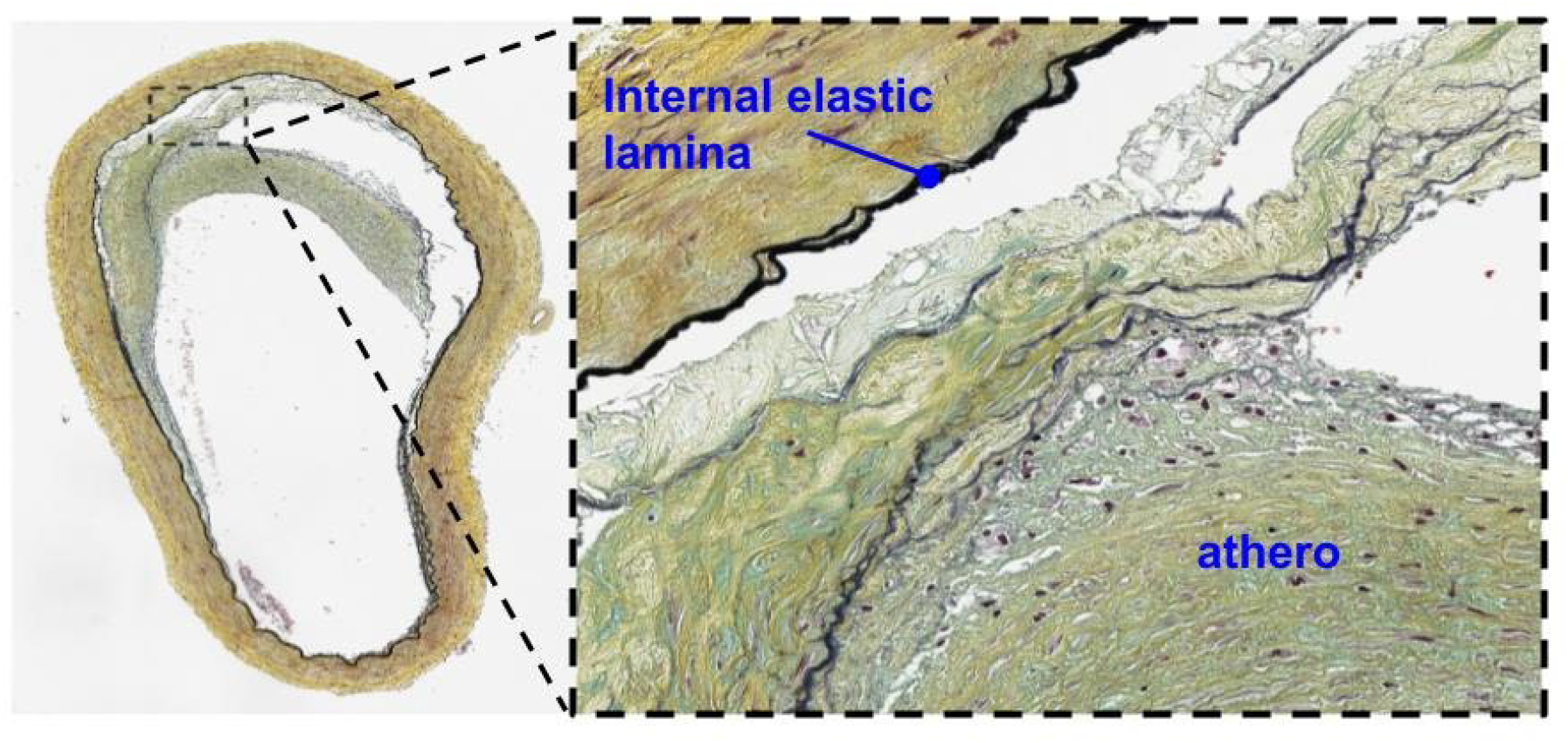
Representative image of formalin-presented right M1. The vessel shows an intact IEL and well-preserved gross morphology. Atherosclerotic plaque is labeled within the vessel lumen.

### Biomechanical Differences between Fresh and Cryopreserved Tissue

Our findings recapitulate the notion that changes in biomechanical properties result from tissue cryopreservation. We demonstrate that cryopreserved vessels are less compliant at both 1.5 and 2.5 PSI than the other experimental groups (Figure 3). Formalin controls boasted the highest strain values over time.

**Figure 3.**
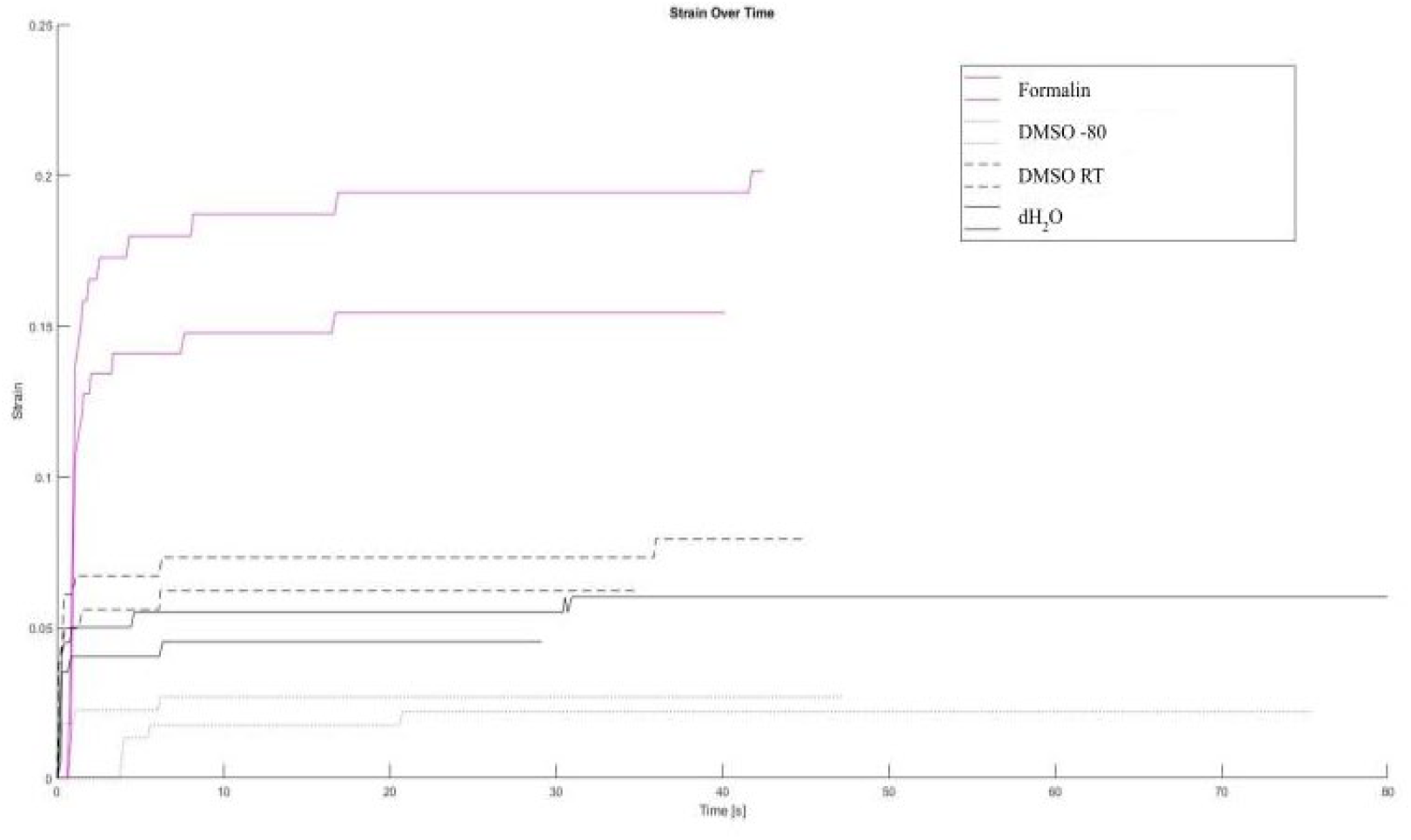
Strain over time curves generated via biaxial inflation testing of right M3 vessels. Formalin controls (solid pink), DMSO −80 °C (dotted), DMSO RT (dashed), and dH_2_O (solid black) are represented. Tandem tests of 1.5 and 2.5 PSI are shown for each experimental group.

### Effects of DMSO Cryopreservation on Cerebrovascular Layer Thicknesses

Counter to prior held suppositions, we found that −80 °C DMSO failed to invariably provide a cryoprotective effect to cerebrovascular tissue. We detected some appreciable differences in vessel tunic thickness. Specifically, the internal elastic lamina (p = 0.03) and adventitia (p < 0.001) thicknesses in MCA territory vessels and the adventitial layers of both ACA (p < 0.001) and SCA (p = 0.04) territories proved inconsistent between experimental groups (Table 4). The total wall thickness of ACA vessels also harbored statistically significant differences between experimental conditions (p < 0.001; Table 4 and Figure S4). Pairwise, within-group comparisons between the dH_2_O, DMSO RT, DMSO −80 °C, and 10% formalin conditions revealed deviations from the formalin-control standard (Table 5). The MCA and ACA saw uncharacteristic elevations in average tissue thickness (p < 0.001 in both cases) in DMSO −80 °C groups compared to formalin controls (Table 5 and Figure S3). This same trend was also seen for the total wall thickness measurements of the ACA (p < 0.001; Table 5 and Figure S4). Aside from the aforementioned elevations in layer thickness in specific DMSO −80 °C experimental groups, the generally observed trend was of no statistically significant nor predictable differences between groups (Table 4 and Figures S1-S4).

**Table 4.**

| Vessel | Vessel Segment Count | p-value | Significance Code |
| --- | --- | --- | --- |
| **Elastin** |  |  |  |
| Middle Cerebral Artery | 296 | 0.03 | * |
| Anterior Cerebral Artery | 141 | 0.08 | ns |
| Posterior Cerebral Artery | 52 | 0.07 | ns |
| Superior Cerebellar Artery | 27 | 0.34 | ns |
| Pericallosal Artery | 26 | 0.27 | ns |
| Internal Carotid Artery | 25 | 0.66 | ns |
| Basilar Artery | 14 | 0.88 | ns |
| Anterior Temporal Artery | 14 | 0.05 | ns |
| **Smooth Muscle** |  |  |  |
| Middle Cerebral Artery | 296 | 0.82 | ns |
| Anterior Cerebral Artery | 141 | 0.15 | ns |
| Posterior Cerebral Artery | 52 | 0.54 | ns |
| Superior Cerebellar Artery | 27 | 0.35 | ns |
| Pericallosal Artery | 26 | 0.44 | ns |
| Internal Carotid Artery | 25 | 0.98 | ns |
| Basilar Artery | 14 | 0.17 | ns |
| Anterior Temporal Artery | 14 | 0.23 | ns |

| **Adventitia** |  |  |  |
| --- | --- | --- | --- |
| Middle Cerebral Artery | 296 | < 0.001 | *** |
| Anterior Cerebral Artery | 141 | < 0.001 | *** |
| Posterior Cerebral Artery | 52 | 0.29 | ns |
| Superior Cerebellar Artery | 27 | 0.04 | * |
| Pericallosal Artery | 26 | 0.53 | ns |
| Internal Carotid Artery | 25 | 0.28 | ns |
| Basilar Artery | 14 | 0.06 | ns |
| Anterior Temporal Artery | 14 | 0.53 | ns |
| **Total Wall** |  |  |  |
| Middle Cerebral Artery | 296 | 0.11 | ns |
| Anterior Cerebral Artery | 141 | < 0.001 | *** |
| Posterior Cerebral Artery | 52 | 0.30 | ns |
| Superior Cerebellar Artery | 27 | 0.36 | ns |
| Pericallosal Artery | 26 | 0.62 | ns |
| Internal Carotid Artery | 25 | 0.37 | ns |
| Basilar Artery | 14 | 0.23 | ns |
| Anterior Temporal Artery | 14 | 0.39 | ns |

**Table 5.**

| **Vessel** | **Pairwise Comparison** | **p-value** | **Significance Code** |
| --- | --- | --- | --- |
| **Elastin** |  |  |  |
| Middle Cerebral Artery | dH2O — DMSO RT | < 0.001 | *** |
|  | dH2O — Formalin | 0.0113 | * |
| **Adventitia** |  |  |  |
| Middle Cerebral Artery | dH2O — Formalin | < 0.001 | *** |
|  | DMSO RT — Formalin | < 0.001 | *** |
|  | DMSO -80 — Formalin | < 0.001 | *** |
| Anterior Cerebral Artery | dH2O — DMSO -80 | 0.01459 | * |
| Anterior Cerebral Artery | DMSO RT — Formalin | < 0.01 | ** |
| Anterior Cerebral Artery | DMSO -80 — Formalin | < 0.001 | *** |
| Superior Cerebellar Artery | dH2O — Formalin | 0.02 | * |
| **Total Wall** |  |  |  |
| Anterior Cerebral Artery | dH2O — Formalin | < 0.01 | ** |
|  | DMSO RT — Formalin | < 0.001 | *** |
|  | DMSO -80 — Formalin | < 0.001 | *** |

### The Effects of DMSO Cryopreservation on Elastin Integrity

In analyzing the effect of −80 °C DMSO cryopreservation on elastin integrity, we observed a paradoxical increase in the number of transverse elastin breaks (Table 6 and Figure 4). This increase was statistically significant in the anterior circulation (p = 0.01), with 5.17 ± 2.86 elastin breaks observed in the −80 °C DMSO group compared to 3.54 ± 2.6 breaks in the formalin control. Further post hoc analyses revealed that, despite not reaching our predetermined threshold for statistical significance, we report a greater number of average elastin breaks in the posterior circulation (5.33 ± 3.14) compared to the anterior circulation (5.17 ± 2.86). Barring discrepancies in sample size due to difficulties with vessel procurement, our findings suggest that the thinner elastin of the posterior circulation may be more vulnerable to fracture following canonical cryopreservation techniques.

**Table 6.**

| **Territory Groups** | **Number of Vessels per Territory Group** | **Number of Segments Scored** | **Average Elastin Breaks per Experimental Group ± SD** |  | **p-value** | **Significance Code** |
| --- | --- | --- | --- | --- | --- | --- |
| Anterior Circulation | 5 | 556 | DMSO -80 | $5.17 \pm 2.86$ | 0.01 | * |
| | | | DMSO RT | $3.71 \pm 2.76$ | | |
| | | | dH<sub>2</sub>O | $3.09 \pm 2.59$ | | |
| | | | Formalin | $3.54 \pm 2.6$ | | |
| Posterior Circulation | 3 | 96 | DMSO -80 | $5.33 \pm 3.14$ | 0.24 | ns |
| | | | DMSO RT | $4.33 \pm 1.97$ | | |
| | | | dH<sub>2</sub>O | $3 \pm 1.9$ | | |
| | | | Formalin | $2.83 \pm 2.14$ | | |

**Figure 4.**
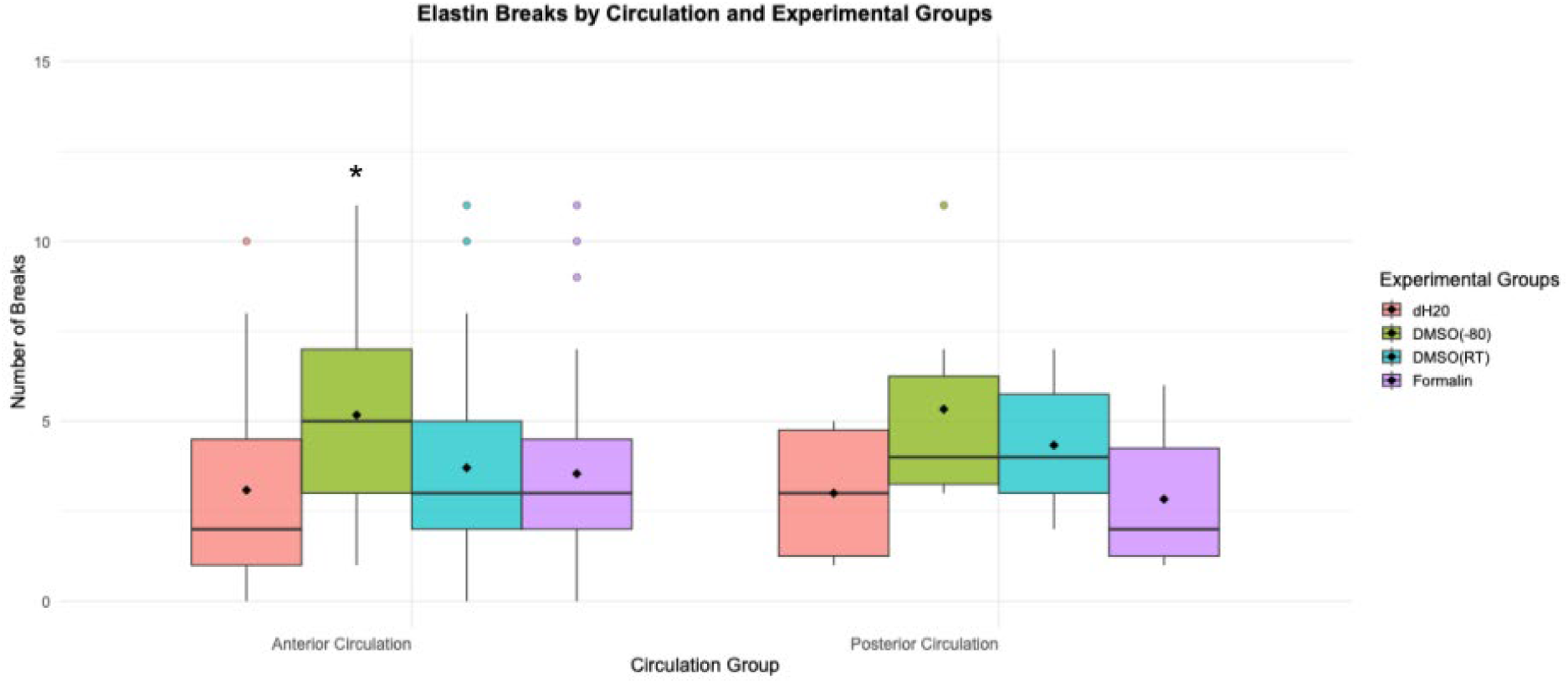
The number of elastin breaks per treatment group is shown, with comparisons between anterior and posterior circulation. Solid lines signify the median and diamonds show the means of each group. Boxes are the inner-quartile range and the whiskers are representative of the maximum and minimum, non-outlier values.

## DISCUSSION

### Biomechanical Testing

In an effort to ensure rigorous study design and maintain precision in the generation of future mechanical testing data, we sought to elucidate whether treatment with cryoprotective agents affected cerebrovascular biomechanics, as the literature fails to reach consensus regarding the degree to which DMSO preserves vascular segments in subzero temperatures [24,25,28,29]. Such analysis had yet to be carried out in smaller-caliber cerebrovascular tissue, thus our study offers a unique perspective. In observing the strain over time relationships between DMSO cryoprotection and mild, biaxial stresses of 1.5 and 2.5 PSI (which roughly approximate a range of physiologic intracranial pressures), we found that −80 °C DMSO storage fails to resemble formalin controls. These data suggest that new approaches to the preservation of cerebrovascular vessels may be warranted.

### Cerebrovascular Layer Thickness

We observed significant alterations in vessel layer thicknesses following −80 °C DMSO storage, particularly in the MCA, ACA, and SCA. In these vessels, MCA IEL and adventitia layers, ACA adventitia and total wall thicknesses, and SCA adventitia deviated significantly from formalin controls. This finding somewhat contrasts the expected findings of cytoprotective agents, as studies of extracranial arteries classically suggest that DMSO-based cryopreservation generally conserves wall thickness [23,28]. For example, human carotid and femoral arteries preserved with DMSO and vapor-phase liquid nitrogen showed no detectable changes in thickness by light or electron microscopy [30]. Similarly, elastomuscular arteries maintained structural integrity and viscoelastic properties across a range of protocols using 5% DMSO, with only minor cryoprotectant-independent delamination noted in rapidly frozen samples [30].

The contrast between extracranial and intracranial findings suggests that cerebral vessels may be uniquely susceptible to cryopreservation-induced alterations. Compared with systemic arteries, intracranial vessels are thinner-walled and possess a higher elastin-to-collagen ratio, which may render them more sensitive to osmotic and freeze/thaw stresses. Even subtle increases in vessel layer construction could affect vascular compliance, redistribution of wall stress, and changes in local hemodynamic response. It is entirely possible that DMSO cryopreservation may induce remodeling of intracranial vessel layers, rather than maintaining baseline histologic architecture. However, the degree to which this occurs across vessel territories seems inconsistent.

### Elastin Integrity

Analysis of elastin continuity revealed an increase in transverse elastin breaks following −80 °C DMSO cryopreservation. This effect was statistically significant in the anterior circulation, where vessels preserved in DMSO exhibited 5.17 ± 2.86 breaks compared with 3.54 ± 2.6 in formalin-fixed controls (p = 0.01). Posterior circulation vessels demonstrated a higher average number of breaks (5.33 ± 3.14) than anterior vessels, though this difference did not reach statistical significance. The increase in elastin discontinuities is consistent with prior reports showing that cryopreservation and thawing exacerbate microfracture formation within the IEL, especially in thinner-walled vessels [29]. The posterior circulation, characterized by a thinner and less structurally robust IEL [17], is likely more susceptible to fragmentation—supporting the concept that baseline wall composition influences preservation-related injury. The presence of IEL microfractures suggests suboptimal preservation and structural compromise, which may manifest in mechanical test data.

## CONCLUSIONS

Our analysis was the first to examine the impact of cryopreservation on the thickness and continuity of human cerebrovascular tissue. Although DMSO is widely used to minimize ice crystal formation during tissue freezing, our findings show that −80 °C DMSO storage did not provide ubiquitous protection. Instead, we observed inconsistent preservation of layer thicknesses and an increase in elastin discontinuities compared to formalin-fixed controls. These alterations suggest that cerebrovascular tissue may be particularly vulnerable to cryopreservation-induced microstructural changes. Our findings suggest that −80 °C DMSO cryopreservation fails to uniformly maintain cerebrovascular tissue architecture and biomechanical properties. Though more testing is needed, this finding prompts the use of fresh tissue when possible. The gold standard for mechanical testing is likely tissue harvested from living or recently deceased individuals, without freezing. Follow-up analyses are needed to confirm these findings and optimize a protocol for the long-term storage of cerebrovascular tissue that conserves vascular identity.

### Limitations

The findings of this study should be interpreted with the following limitations in mind. First, the data presented herein is dominated by samples collected from the anterior circulation, as posterior vessels are more difficult to resect. Future investigations should include equal representation of anterior and posterior vessels. Second, there exists a degree of subjectivity when scoring vessel thicknesses. Despite attempts to correct for individual bias by incorporating two blinded reviewers, slight differences in layer interpretation does occur and may account for small deviations in the data presented.

## SUPPLEMENTAL FIGURES

**Figure S1.**
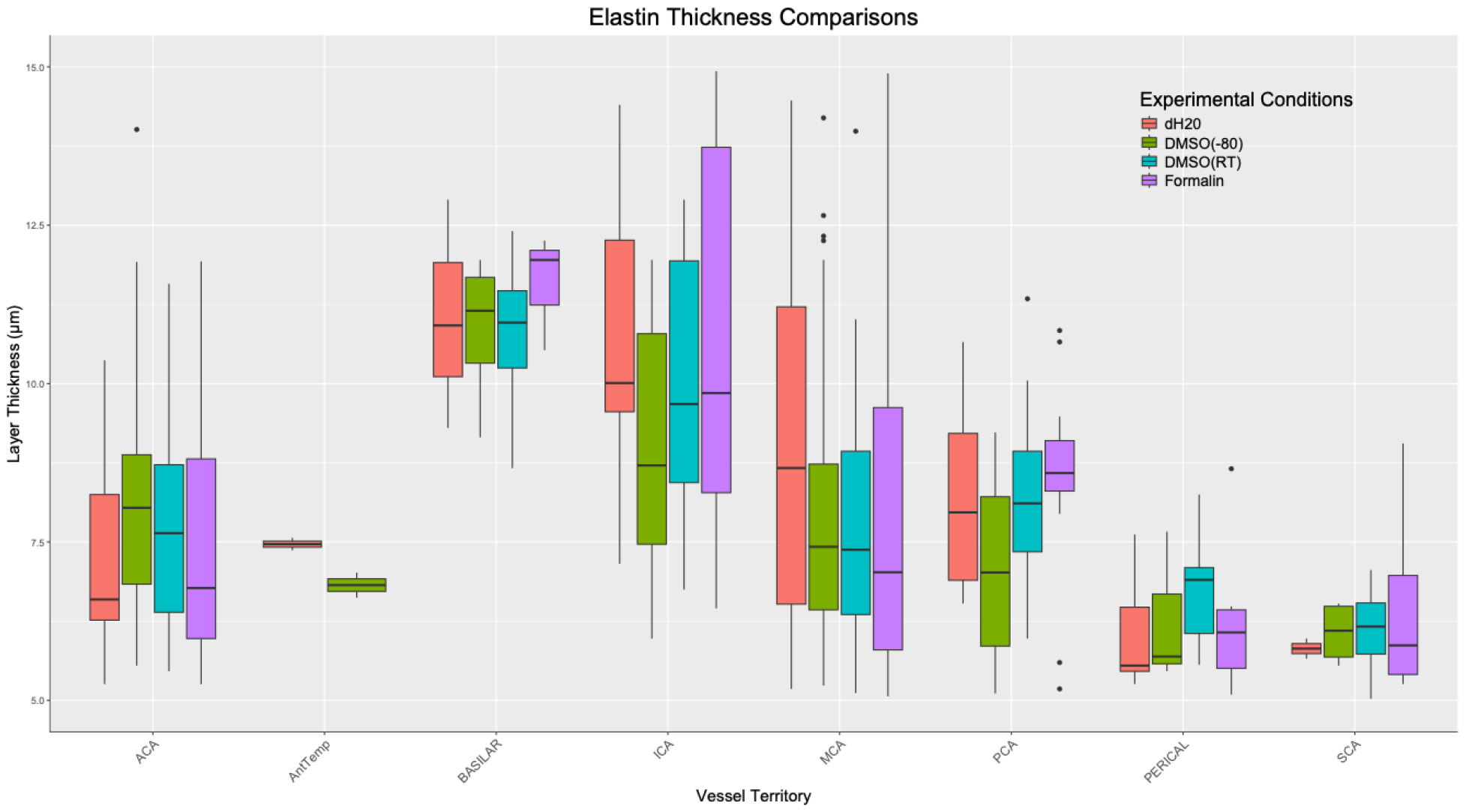
Elastin thickness comparisons of the vessel territories represented in this study, inclusive of formalin controls, DMSO −80 °C, DMSO RT, and dH_2_O experimental conditions. Solid lines signify the median and diamonds show the means of each group. Boxes are the inner-quartile range and the whiskers are representative of the maximum and minimum, non-outlier values.

**Figure S2.**
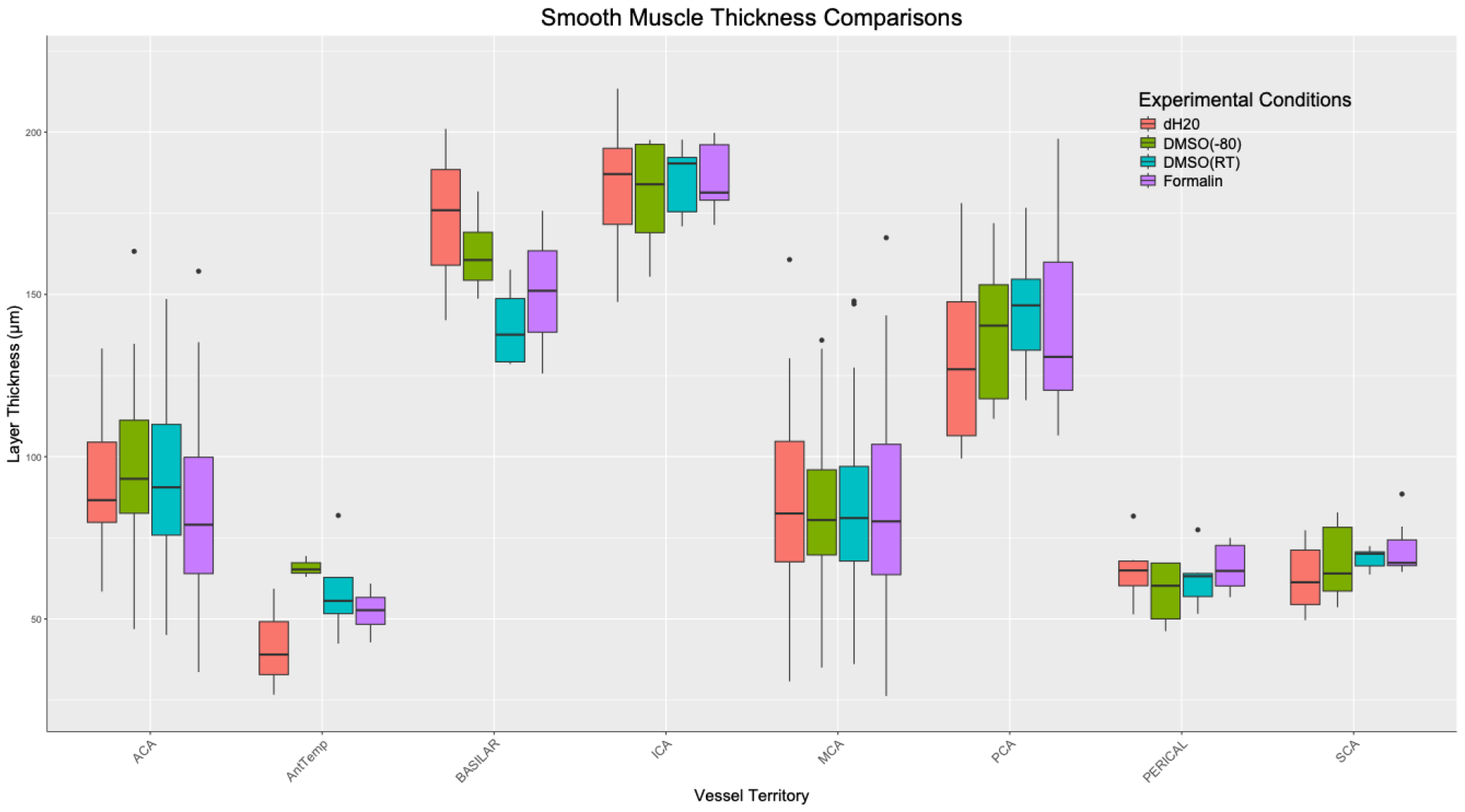
Smooth muscle thickness comparisons of the vessel territories represented in this study, inclusive of formalin controls, DMSO −80 °C, DMSO RT, and dH_2_O experimental conditions. Solid lines signify the median and diamonds show the means of each group. Boxes are the inner-quartile range and the whiskers are representative of the maximum and minimum, non-outlier values.

**Figure S3.**
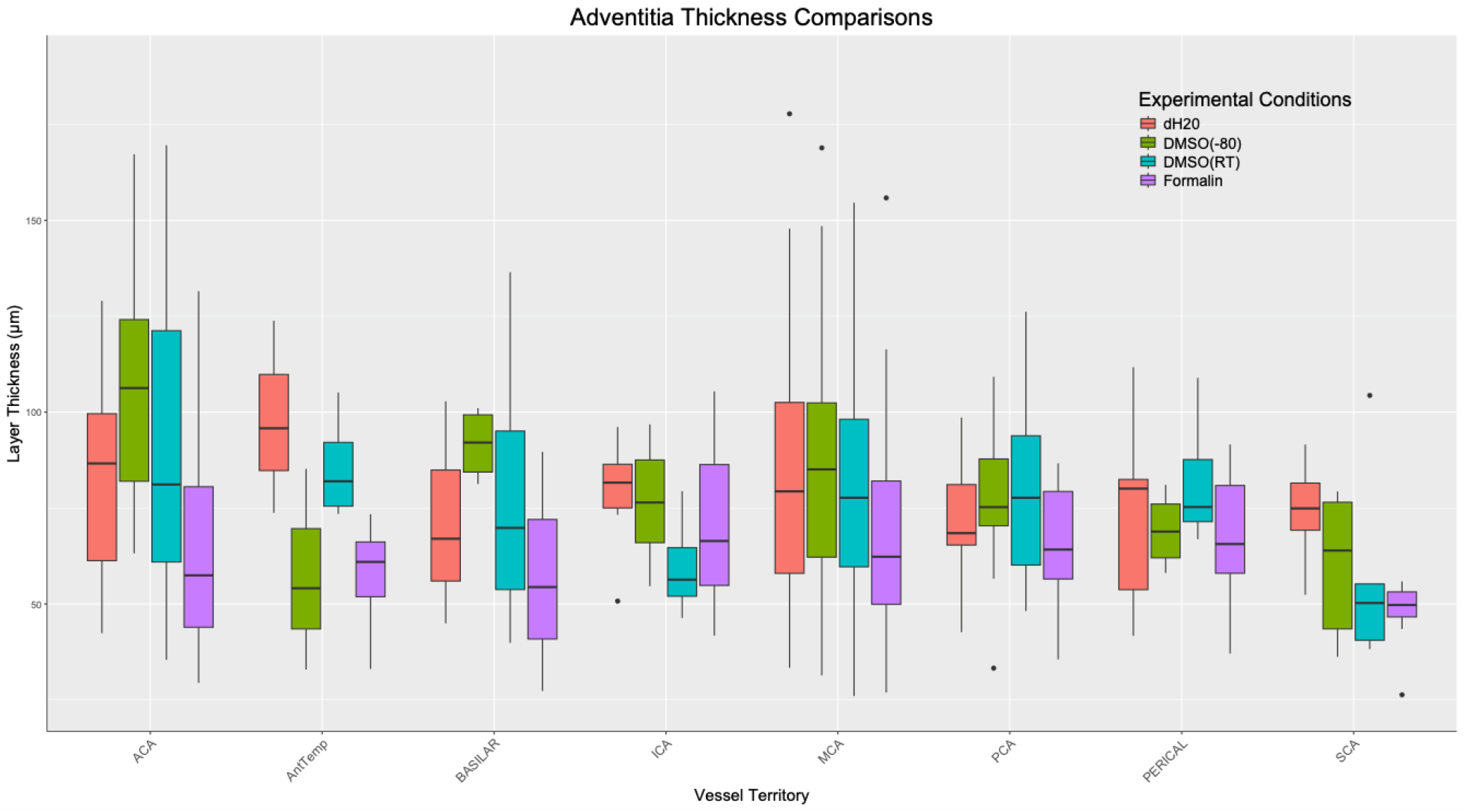
Adventitia thickness comparisons of the vessel territories represented in this study, inclusive of formalin controls, DMSO −80 °C, DMSO RT, and dH_2_O experimental conditions. Solid lines signify the median and diamonds show the means of each group. Boxes are the inner-quartile range and the whiskers are representative of the maximum and minimum, non-outlier values.

**Figure S4.**
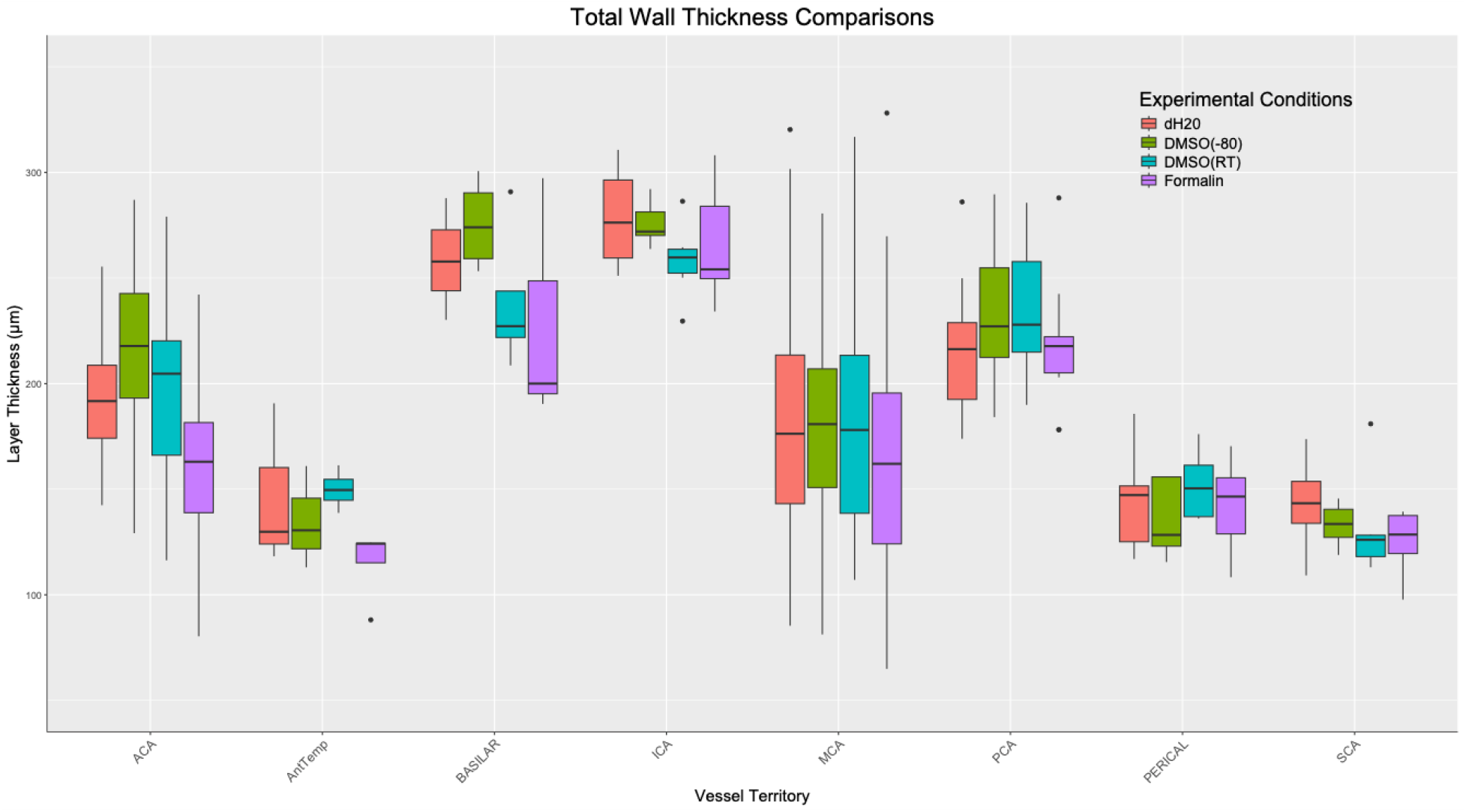
Total wall thickness comparisons of the vessel territories represented in this study, inclusive of formalin controls, DMSO −80 °C, DMSO RT, and dH_2_O experimental conditions. Solid lines signify the median and diamonds show the means of each group. Boxes are the inner-quartile range and the whiskers are representative of the maximum and minimum, non-outlier values.

## Notes

### Competing Interest Statement

The authors have declared no competing interest.

## REFERENCES

1. Roth GA, Mensah GA, Johnson CO, Addolorato G, Ammirati E, Baddour LM, et al. Global Burden of Cardiovascular Diseases and Risk Factors, 1990-2019: Update From the GBD 2019 Study. J Am Coll Cardiol. 2020;76: 2982–3021. doi:10.1016/j.jacc.2020.11.010

2. GBD 2015 Neurological Disorders Collaborator Group. Global, regional, and national burden of neurological disorders during 1990-2015: a systematic analysis for the Global Burden of Disease Study 2015. Lancet Neurol. 2017;16: 877–897. doi:10.1016/S1474-4422(17)30299-5

3. GBD 2021 Stroke Risk Factor Collaborators. Global, regional, and national burden of stroke and its risk factors, 1990-2021: a systematic analysis for the Global Burden of Disease Study 2021. Lancet Neurol. 2024;23: 973–1003. doi:10.1016/S1474-4422(24)00369-7

4. Weber R, Eyding J, Kitzrow M, Bartig D, Weimar C, Hacke W, et al. Distribution and evolution of acute interventional ischemic stroke treatment in Germany from 2010 to 2016. Neurol Res Pract. 2019;1: 4. doi:10.1186/s42466-019-0010-8

5. Atchaneeyasakul K, Liaw N, Lee RH, Liebeskind DS, Saver JL. Patterns of mechanical thrombectomy for stroke before and after the 2015 pivotal trials and US national guideline update. J Stroke Cerebrovasc Dis. 2020;29: 105292. doi:10.1016/j.jstrokecerebrovasdis.2020.105292

6. Nogueira RG, Jadhav AP, Haussen DC, Bonafe A, Budzik RF, Bhuva P, et al. Thrombectomy 6 to 24 Hours after Stroke with a Mismatch between Deficit and Infarct. N Engl J Med. 2018;378: 11–21. doi:10.1056/NEJMoa1706442

7. Etminan N, Chang H-S, Hackenberg K, de Rooij NK, Vergouwen MDI, Rinkel GJE, et al. Worldwide Incidence of Aneurysmal Subarachnoid Hemorrhage According to Region, Time Period, Blood Pressure, and Smoking Prevalence in the Population: A Systematic Review and Meta-analysis. JAMA Neurol. 2019;76: 588–597. doi:10.1001/jamaneurol.2019.0006

8. Berman MF, Sciacca RR, Pile-Spellman J, Stapf C, Connolly ES, Mohr JP, et al. The epidemiology of brain arteriovenous malformations. Neurosurgery. 2000;47: 389–96; discussion 397. doi:10.1097/00006123-200008000-00023

9. Al-Shahi R, Warlow C. A systematic review of the frequency and prognosis of arteriovenous malformations of the brain in adults. Brain. 2001;124: 1900–1926. doi:10.1093/brain/124.10.1900

10. Kondziolka D, McLaughlin MR, Kestle JR. Simple risk predictions for arteriovenous malformation hemorrhage. Neurosurgery. 1995;37: 851–855. doi:10.1227/00006123-199511000-00001

11. Karlsson B, Jokura H, Yang H-C, Yamamoto M, Martinez R, Kawagishi J, et al. Clinical outcome following cerebral AVM hemorrhage. Acta Neurochir (Wien). 2020;162: 1759–1766. doi:10.1007/s00701-020-04380-z

12. Rutledge C, Cooke DL, Hetts SW, Abla AA. Brain arteriovenous malformations. Handb Clin Neurol. 2021;176: 171–178. doi:10.1016/B978-0-444-64034-5.00020-1

13. Mohr JP, Parides MK, Stapf C, Moquete E, Moy CS, Overbey JR, et al. Medical management with or without interventional therapy for unruptured brain arteriovenous malformations (ARUBA): a multicentre, non-blinded, randomised trial. Lancet. 2014;383: 614–621. doi:10.1016/S0140-6736(13)62302-8

14. Luther E, McCarthy DJ, Burks J, Govindarajan V, Lu VM, Silva M, et al. National reduction in cerebral arteriovenous malformation treatment correlated with increased rupture incidence. J Neurointerv Surg. 2022; neurintsurg-2022-019110. doi:10.1136/jnis-2022-019110

15. Laurence DW, Homburg H, Yan F, Tang Q, Fung K-M, Bohnstedt BN, et al. A pilot study on biaxial mechanical, collagen microstructural, and morphological characterizations of a resected human intracranial aneurysm tissue. Sci Rep. 2021;11: 3525. doi:10.1038/s41598-021-82991-x

16. Hill MA, Nourian Z, Ho I-L, Clifford PS, Martinez-Lemus L, Meininger GA. Small Artery Elastin Distribution and Architecture-Focus on Three Dimensional Organization. Microcirculation. 2016;23: 614–620. doi:10.1111/micc.12294

17. Thiyagarajah N, Witek A, Davison M, Butler R, Erdemir A, Tsiang J, et al. Histological analysis of intracranial cerebral arteries for elastin thickness, wall thickness, and vessel diameters: an atlas for computational modeling and a proposed predictive multivariable model of elastin thickness. J Clin Med. 2025;14. doi:10.3390/jcm14124320

18. Macrae RA, Miller K, Doyle BJ. Methods in mechanical testing of arterial tissue: A review. Strain. 2016;52: 380–399. doi:10.1111/str.12183

19. Notman R, Noro M, O’Malley B, Anwar J. Molecular basis for dimethylsulfoxide (DMSO) action on lipid membranes. J Am Chem Soc. 2006;128: 13982–13983. doi:10.1021/ja063363t

20. Lee E, Baiz CR. How cryoprotectants work: hydrogen-bonding in low-temperature vitrified solutions. Chem Sci. 2022;13: 9980–9984. doi:10.1039/d2sc03188d

21. Akhoondi M, Oldenhof H, Sieme H, Wolkers WF. Freezing-induced cellular and membrane dehydration in the presence of cryoprotective agents. Mol Membr Biol. 2012;29: 197–206. doi:10.3109/09687688.2012.699106

22. Rendal E, Santos MVM, Rodriguez M, Sánchez J, Segura R, Matheu G, et al. Effects of cryopreservation and thawing on the structure of vascular segment. Transplant Proc. 2004;36: 3283–3287. doi:10.1016/j.transproceed.2004.10.057

23. Pascual G, García-Honduvilla N, Rodríguez M, Turégano F, Bujan J, Bellón JM. Effect of the thawing process on cryopreserved arteries. Ann Vasc Surg. 2001;15: 619–627. doi:10.1007/s100160010130

24. Venkatasubramanian RT, Grassl ED, Barocas VH, Lafontaine D, Bischof JC. Effects of freezing and cryopreservation on the mechanical properties of arteries. Ann Biomed Eng. 2006;34: 823–832. doi:10.1007/s10439-005-9044-x

25. Venkatasubramanian RT, Wolkers WF, Shenoi MM, Barocas VH, Lafontaine D, Soule CL, et al. Freeze-thaw induced biomechanical changes in arteries: role of collagen matrix and smooth muscle cells. Ann Biomed Eng. 2010;38: 694–706. doi:10.1007/s10439-010-9921-9

26. Fonck E, Feigl GG, Fasel J, Sage D, Unser M, Rüfenacht DA, et al. Effect of aging on elastin functionality in human cerebral arteries. Stroke. 2009;40: 2552–2556. doi:10.1161/STROKEAHA.108.528091

27. Schindelin J, Arganda-Carreras I, Frise E, Kaynig V, Longair M, Pietzsch T, et al. Fiji: an open-source platform for biological-image analysis. Nat Methods. 2012;9: 676–682. doi:10.1038/nmeth.2019

28. Kostelnik CJ, Crouse KJ, Goldsmith JD, Eberth JF. Impact of cryopreservation on elastomuscular artery mechanics. J Mech Behav Biomed Mater. 2024;154: 106503. doi:10.1016/j.jmbbm.2024.106503

29. Buján J, Pascual G, García-Honduvilla N, Gimeno MJ, Jurado F, Carrera-San Martín A, et al. Rapid thawing increases the fragility of the cryopreserved arterial wall. Eur J Vasc Endovasc Surg. 2000;20: 13–20. doi:10.1053/ejvs.2000.1090

30. Rosset E, Friggi A, Novakovitch G, Rolland PH, Rieu R, Pellissier JF, et al. Effects of cryopreservation on the viscoelastic properties of human arteries. Ann Vasc Surg. 1996;10: 262–272. doi:10.1007/BF02001892

